# Astrocyte coverage of excitatory synapses correlates to measures of synapse structure and function in primary visual cortex

**DOI:** 10.1101/2023.12.01.569664

**Authors:** Connon I. Thomas, Melissa A. Ryan, Micaiah C. McNabb, Naomi Kamasawa, Benjamin Scholl

## Abstract

Most excitatory synapses in the mammalian brain are contacted by astrocytes, forming the tripartite synapse. This interface is thought to be critical for glutamate turnover and structural or functional dynamics of synapses. While the degree of synaptic contact of astrocytes is known to vary across brain regions and animal species, the implications of this variability remain unknown. Furthermore, precisely how astrocyte coverage of synapses relates to *in vivo* functional properties of individual dendritic spines has yet to be investigated. Here, we characterized perisynaptic astrocyte processes (PAPs) contacting synapses of pyramidal neurons of the ferret visual cortex and, using correlative light and electron microscopy, examined their relationship to synaptic strength and to sensory-evoked Ca^2+^ activity. Nearly all synapses were contacted by PAPs, and most were contacted along the axon-spine interface (ASI). Structurally, we found that the degree of PAP coverage scaled with synapse size and complexity. Functionally, we found that PAP coverage scaled with the selectivity of Ca^2+^ responses of individual synapses to visual stimuli and, at least for the largest synapses, scaled with the reliability of visual stimuli to evoke postsynaptic Ca^2+^ events. Our study shows astrocyte coverage is highly correlated with structural properties of excitatory synapses in the visual cortex and implicates astrocytes as a contributor to reliable sensory activation.

## Introduction

Excitatory glutamatergic synapses comprise the majority of synaptic connections in the central nervous system. Fine protrusions from astrocytes commonly surround pre- and postsynaptic compartments of excitatory synapses, forming the tripartite synapse and allowing for a wide range of molecular interactions (Semyanov and Verkhratsky, 2021; Aten et al., 2022). These perisynaptic astrocyte processes (PAPs) are implicated in de novo glutamate synthesis from glucose (Hertz et al., 1999; Marx et al., 2015), synapse formation and pruning (Allen et al., 2012; Tsai et al., 2012; Chung et al., 2013; Singh et al., 2016; Stogsdill et al., 2017; Sipe et al., 2021), gliotransmitter release (Blanco-Suarez et al., 2018), glutamate buffering and recycling (Marx et al., 2015; Rose et al., 2018; Torres-Ceja and Olsen, 2022), maturation and modulation of synaptic strength (Jourdain et al., 2007; Nishida and Okabe, 2007; Henneberger et al., 2010; Jones et al., 2011), and provision of metabolites to neurons (Bélanger et al., 2011; Suzuki et al., 2011; Bonvento and Bolaños, 2021). This variety of roles attributed to astrocytes identifies them as an important and complex player in the maintenance and function of synapses. Synapses are similarly diverse, exhibiting heterogeneity in morphology (Svoboda and Holtmaat, 2009), receptor expression (Favuzzi and Rico, 2018), and presynaptic input drive (Scholl and Fitzpatrick, 2020). However, precisely how PAPs support synaptic diversity is still unclear.

Synaptic coverage by PAPs varies across brain regions and neural circuits, with no exact consensus on their relationship with individual synapses. Structural measurements in the murine hippocampus and amygdala suggest no correlation between synapse size and astrocyte coverage length at the axon-spine interface (ASI) (Witcher et al., 2007; Ostroff et al., 2014). This led to a proposal that large synapses have proportionally less coverage and are more susceptible to glutamate spillover from synaptic clefts (Witcher et al., 2007; Herde et al., 2020), but other evidence suggests that small synapses have a greater degree of glutamate spillover (Gavrilov et al., 2018). In barrel cortex, synapse size and PAP coverage length are reported to be correlated (Genoud et al., 2006). Cortical synapse-astrocyte coverage also appears to vary by layer, such that ensheathment of the synapse is greater in layer 2/3 than layer 6 (Kasthuri et al., 2015; Lanjakornsiripan et al., 2018). Simply put, the relationship between PAPs and synapse morphology remains puzzling. Even less is known about the relationship between PAP coverage and synaptic activity, as investigation requires a challenging combination of high resolution structural imaging and in vivo imaging. PAP coverage may follow a universal principle yet to be elucidated or depend on brain region, functional specialization, neuronal cell type, contact site, or animal species. Given the structural and functional heterogeneity of synapses, it is reasonable to hypothesize that astrocytic coverage is neither random nor uniform; yet, no direct measurements have been made relating astrocyte coverage to structure and function of individual dendritic spines.

To advance our understanding of tripartite synapses, correlative studies provide a means to integrate structural and functional details (Papouin et al., 2017; Zhou et al., 2019; Semyanov and Verkhratsky, 2021; Torres-Ceja and Olsen, 2022). Following this vein, we examined a correlative light and electron microscopy (CLEM) dataset (Scholl et al., 2021) and volumetrically reconstructed astrocytic processes surrounding synapses onto dendritic spines of layer 2/3 pyramidal neurons of ferret primary visual cortex (V1). Using 3D reconstructions from Serial Block-Face Scanning Electron Microscopy (SBF-SEM), we measured PAP coverage of the pre- and postsynaptic membrane compartments (i.e. the synaptic assembly) as well as the coverage of the ASI. We then compared coverage to synapse structural characteristics and *in vivo* visually-evoked Ca^2+^ activity of previously imaged dendritic spines. Almost all dendritic spines on basal dendrites of layer 2/3 excitatory neurons in ferret V1 were contacted by astrocytes (99% of synaptic assemblies). In contrast to some previous studies, PAP coverage area and length scaled with synapse size, while the fraction of coverage was independent. In addition, synapses lacking PAP coverage were significantly smaller than those that had at least some coverage, and complex synapses had greater coverage. When comparing Ca^2+^ activity evoked by visual stimuli, we found PAP coverage was correlated with tuning selectivity and trial-to-trial response reliability. Our results, the first of their kind, provide direct evidence that astrocytic coverage is correlated with synapse size and functional activity. We hypothesize that the high degree of astrocyte coverage in ferret V1 is a hallmark of sensory cortex and supports synaptic reliability of strong synapses through enhanced glutamate turnover.

## Methods

All procedures were performed according to NIH guidelines and approved by the Institutional Animal Care and Use Committee at Max Planck Florida Institute for Neuroscience.

### Animals, Viral Injections, and Cranial Windows

Information on animals, survival viral injections, cranial window implantation are described in depth in our previous studies (Scholl et al., 2017, 2019, 2021; Thomas et al., 2023). Briefly, layer 2/3 neurons of the primary visual cortex of female ferrets (n = 3, Marshall Farms) were driven to express the Ca^2+^ indicator GCaMP6s via a Cre-dependent AAV expression system. On the day of imaging, a cranial window was surgically implanted to image Ca^2+^ activity of cortical pyramidal neurons during presentation of visual stimuli.

### Two-Photon Imaging

Two-photon imaging was performed using a Bergamo II microscope (Thorlabs) running Scanimage (Vidrio Technologies) with 940 nm dispersion-compensated excitation provided by an Insight DS+ (Spectraphysics). For spine and axon imaging, power after the objective was limited to <50 mW. Images were collected at 30 Hz using bidirectional scanning with 512 × 512 pixel resolution or with custom ROIs (region of interest; framerate range: 22 - 50 Hz). Somatic imaging was performed with a resolution of 0.488 - 0.098 µm/pixel. Dendritic spine imaging was performed with a resolution of 0.164 - 0.065 µm/pixel.

### Visual Stimuli

Visual stimuli were generated using Psychopy (Peirce, 2007). The monitor was placed 25 cm from the animal. Receptive field locations for each cell were hand mapped and the spatial frequency optimized (range: 0.04 to 0.20 cpd). For each soma and dendritic segment, square-wave or sine-wave drifting gratings were presented at 22.5 degree increments to each eye independently (2 second duration, 1 second ISI, 8-10 trials for each field of view). Drifting gratings of different directions (0 – 315°) were presented independently to both eyes with a temporal frequency of 4 Hz.

### Two-Photon Imaging Analysis

Imaging data were excluded from analysis if motion along the z-axis was detected. Dendrite images were corrected for in-plane motion via a 2D cross-correlation based approach in MATLAB or using a piecewise non-rigid motion correction algorithm (Pnevmatikakis and Giovannucci, 2017). ROIs were drawn in ImageJ; dendritic ROIs spanned contiguous dendritic segments and spine ROIs were fit with custom software. Mean pixel values for ROIs were computed over the imaging time series and imported into MATLAB. ΔF/F was computed with a time-averaged median or percentile filter (10th percentile). We subtracted a scaled version of the dendritic signal to remove back-propagating action potentials as performed previously (Wilson et al., 2016). ΔF/F traces were synchronized to stimulus triggers sent from Psychopy and collected by Spike2.

Peak responses to bars and gratings were computed using the Fourier analysis to calculate mean and modulation amplitudes for each stimulus presentation, which were summed together. Spines were included for analysis if the mean peak for the preferred stimulus was >10%, the SNR at the preferred stimulus was > 1, and spines were weakly correlated with the dendritic signal (Spearman’s correlation, r < 0.4). Some spine traces contained negative events after subtraction, so correlations were computed ignoring negative values. Preferred orientation for each spine was calculated by fitting responses with a double Gaussian tuning curve (Wilson et al., 2016) using lsqcurvefit (Matlab). Orientation selectivity was computed by calculating the vector strength of mean responses (Wilson et al., 2016). For each tuned spine (orientation selectivity > 0.1), we calculated orientation preference difference between the spine and parent soma as in Wilson et al. (2016). To calculate response reliability, we converted spine Ca^2+^ responses on individual trials to binary events: single trial responses were filtered with exponential kernel, the response peak amplitude was identified, and ‘events ‘were defined as peaks exceeding 2 standard deviations above the ΔF/F noise of each spine. To compute mean reliability, we averaged across stimuli evoking an average response greater than 10% ΔF/F.

### Perfusion, fixation, and slice preparation for fluorescence imaging

Anesthetized animals were immediately perfused with 2% paraformaldehyde and 2 – 2.5% glutaraldehyde in a 0.1 M sodium cacodylate buffer (pH 7.4). Following removal, brains were sliced at 80 µm parallel to the area flattened by the cranial window. Slices were quickly imaged at low magnification (20x, 0.848 × 0.848 µm/pixel) using a Leica CLSM TCS SP5 II running LAS AF (ver. 3.0, Leica) with 488 nm laser excitation. Fluorescence of GCaMP6s was used to locate the target cell in this view.

Autofluorescence resulting from glutaraldehyde fixation also produced signal within the tissue slices. A field of view large enough to cover roughly one-fourth of the slice, with the target cell included, was captured using image tiling. This process was performed to identify the fluorescent cell of interest and for slice-level correlation in later steps of the workflow. The slice containing the target cell was then imaged with the same 20x objective at higher pixel resolution (0.360 × 0.360 × 0.976 µm/voxel) to obtain a z-stack of the full depth of the slice and immediate region surrounding the target cell. CLSM imaging required approximately 1-3 hours to find the cell of interest within a slice and capture a tiled slice overview and higher resolution z-stack.

### Sample preparation for SBF-SEM imaging

Samples were prepared as previously described (Scholl et al., 2021; Thomas et al., 2021). Briefly, tissue pieces were incubated in an aqueous solution of 2% osmium tetroxide buffered in 0.1 M sodium cacodylate for 45 minutes at room temperature (RT). Tissue was not rinsed and the osmium solution was replaced with cacodylate buffered 2.5% potassium ferrocyanide for 45 minutes at RT in the dark. Tissue was rinsed with water 2 × 10 minutes, which was repeated between consecutive steps. Tissue was incubated at room temperature for 20 minutes in aqueous 1% thiocarbohydrizide dissolved at 60°C, aqueous 1% osmium tetroxide for 45 minutes at RT, and then 1% uranyl acetate in 25% ethanol for 20 minutes at RT in the dark. Tissue was rinsed then left in water overnight at 4°C. The following day, tissue was stained with Walton’s lead aspartate for 30 minutes at 60°C. Tissue was then dehydrated in a graded ethanol series (30, 50, 70, 90, 100%), 1:1 ethanol to acetone, then 100% dry acetone. Tissue was infiltrated using 3:1 acetone to Durcupan resin (Sigma Aldrich) for 2 hours, 1:1 acetone to resin for 2 hours, and 1:3 acetone to resin overnight, then flat embedded in 100% resin on a glass slide and covered with an Aclar sheet at 60°C for 2 days. The tissue was trimmed to less than 1 × 1 mm in size, the empty resin at the surface was shaved to expose tissue surface using an ultramicrotome (UC7, Leica), then turned downwards to be remounted to a metal pin with conductive silver epoxy (CircuitWorks, Chemtronics).

### SBF-SEM image acquisition and volume data handling

Samples were imaged using a correlative approach as previously described (Thomas et al., 2021). Briefly, tissue was sectioned and imaged using 3View2XP and Digital Micrograph (ver. 3.30.1909.0, Gatan Microscopy Suite) installed on a Gemini SEM300 (Carl Zeiss Microscopy LLC.) equipped with an OnPoint BSE detector (Gatan, Inc.). The detector magnification was calibrated within SmartSEM imaging software (ver. 6.0, Carl Zeiss Microscopy LLC.) and Digital Micrograph with a 500 nm cross line grating standard. A low magnification image of each block face was manually matched to its corresponding depth in the CLSM Z-stack using blood vessels and cell bodies as fiducials. Final imaging was performed at 2.0 - 2.2 kV accelerating voltage, 20 or 30 µm aperture, working distance of ∼5 mm, 0.5 - 1.2 µs pixel dwell time, 5.7 - 7 nm per pixel, knife speed of 0.1 mm/sec with oscillation, and 56 - 84 nm section thickness. Acquisition was automated and ranged from several days to several weeks depending on the size of the ROI and imaging conditions. The true section thickness was measured using mitochondria diameter calibrations (Fiala and Harris, 2001). Calibration for each block was required, since variation in thickness can occur due to heating, charging from the electron beam, resin polymerization, and tissue fixation and staining quality (Starborg et al., 2013; Hughes et al., 2014).

Serial tiled images were exported as TIFFs to TrakEM2 (Schindelin et al., 2012) within ImageJ (ver. 1.52p) to montage tiles, then aligned using Scale-Invariant Feature Transform image alignment with linear feature correspondences and rigid transformation (Lowe, 2004). Once aligned, images were inverted and contrast normalized.

### SBF-SEM image analysis

Aligned images were exported to Microscopy Image Browser (ver. 2.51, 2.6, Belevich et al., 2016) for segmentation of dendrites, spines, postsynaptic densities (PSDs), and boutons. Three annotators preformed segmentation and the segmentations of each annotator were proof-read by an experienced annotator (∼1000 hours of segmentation experience) prior to quantification. Binary labels files were imported to Amira (ver. 6.7, 2019.1, Thermo Fisher Scientific) which was used to create 3D surface models of each dendrite, spine, and PSD. Once reconstructed, the model of each dendrite was manually overlaid onto its corresponding two-photon image using Adobe Photoshop for re-identification of individual spines. Amira was used to measure the volume of spine heads, surface area of PSDs, and spine neck length.

For measurements of astrocyte coverage of the synaptic assembly (i.e. pre-/postsynaptic compartment) membrane, portions of astrocyte that were in contact with the membranes of spines and boutons were segmented in 3D within Microscopy Image Browser. Amira was then used to generate meshes and measure the apposed surface area on spines and boutons. To measure ASI-PAP coverage, the same meshes generated using Amira were exported to the virtual reality program syGlass (ver. 1.7.2). The ASI was measured manually in a 3D space as the perimeter of the spine and bouton apposition. The length of the astrocyte contact was then measured as the line along that perimeter where the astrocyte, spine, and bouton meet (**Figure 1E**). Spines, boutons, and astrocytes were defined to be in apposition if their membranes were less than 50 nm apart in space. We defined a “discrete astrocyte segment” as a segment that contacts the ASI, moves more than 50 nm away, then approaches the ASI again.

**Figure 1:**
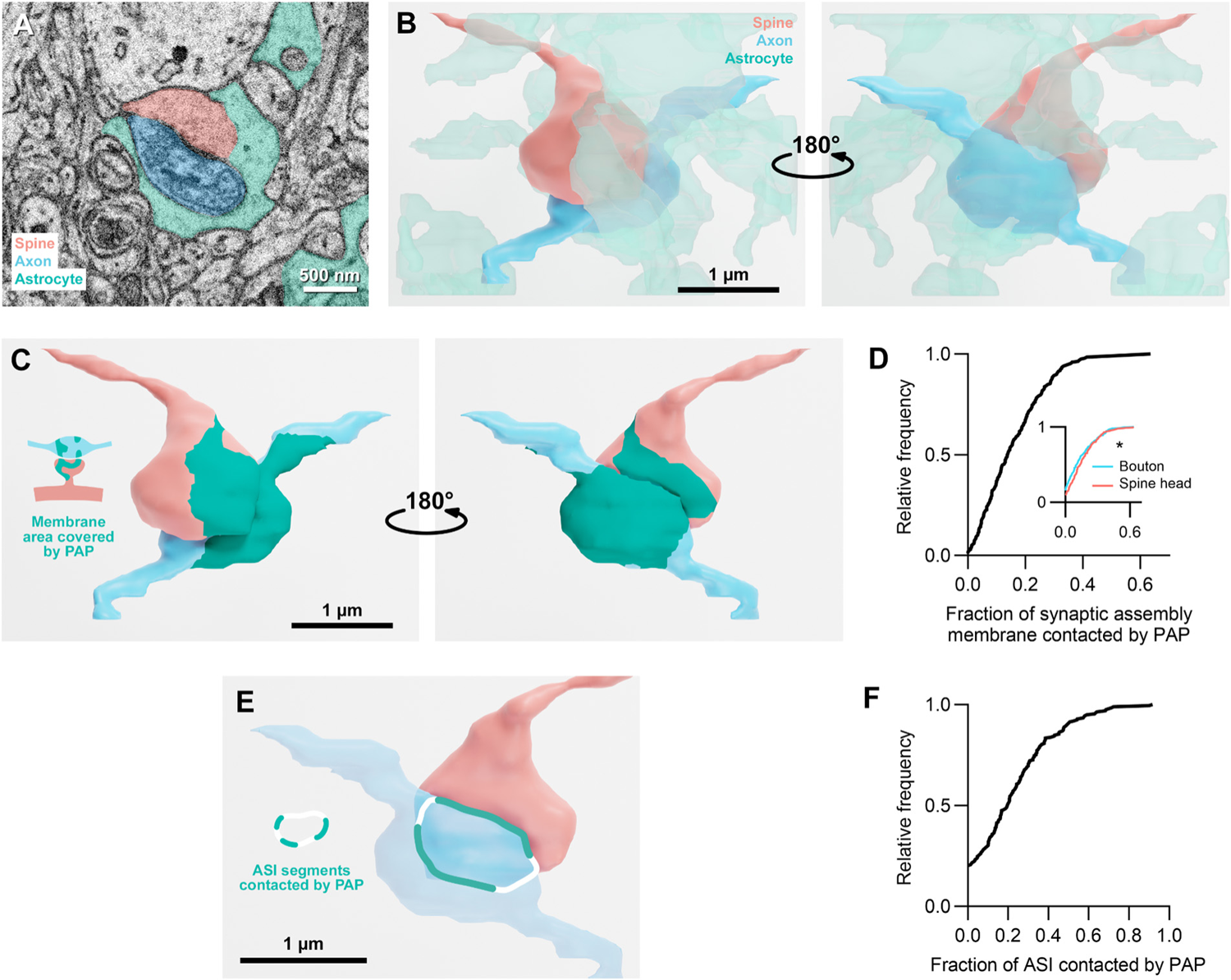
Astrocyte coverage of the synaptic assembly membrane and axon-spine interface. **A.** SEM micrograph showing a PAP surrounding a synapse. **B.** 3D volume rendering of the same spine and axon in A, with astrocyte processes present in the volume (within a 4 µm radius cylinder) rendered transparent. **C**. 3D rendering and illustration of membrane area covered by the PAP. **D.** Cumulative distribution function (CDF) of the fraction of combined spine head and bouton surface contacted by the PAP. **E.** 3D rendering and illustration of the ASI segments contacted by astrocyte. **F.** CDF of the fraction of ASI contacted by PAP. *p < 0.05.

### Statistics

Statistical analyses are described in the main text and in figure legends. We used non-parametric statistical analyses including the Wilcoxon sign-rank test, Mann-Whitney rank-sum test (MW), Kruskal-Wallis test (KW), or permutation tests to avoid assumptions about the distributions of the data. Statistical analysis was performed in MATLAB (2021b) or GraphPad Prism (ver. 9). Circular correlation coefficients were computed for tests with circular variables. For all other tests, Spearman’s correlation coefficient was computed. All correlation significance tests were one-sided. Synapses with no coverage by astrocyte were excluded from relevant correlations.

## Results

### Astrocytes contact nearly all excitatory synapses on basal dendrites of ferret V1 neurons

Astrocyte processes were identified by their complex shape and the presence of dark glycogen granules in their lucent cytoplasm (Spacek, 1985; **Figure 1A**). We targeted spine and bouton reconstructions from our previously published volume EM dataset of excitatory synapses on basal dendrites of layer 2/3 pyramidal neurons in ferret V1 (Scholl et al., 2021), manually segmenting astrocyte processes surrounding each synapse (**Figure 1B**; see Methods). We first quantified the area of pre- and postsynaptic membrane contacted by a PAP (**Figure 1C**) due to its importance as a site of glutamine and lactate transfer between astrocytes and neurons (Bröer and Brookes, 2001; Verkhratsky et al., 2015). We found that 99% of synaptic assemblies examined are contacted by an astrocyte on either the spine or bouton membrane (n = 196/198; **Figure 1D**). Median coverage over the entire synapse assembly was 14%, with a maximum of 64%. As a previous study in rat CA1 and cerebellum found PAPs are more likely to provide postsynaptic coverage (Lehre and Rusakov, 2002), we considered spine head and bouton coverage individually. Spine heads had a slightly higher median percentage of contact (median spine head membrane coverage percentage = 13.5%, IQR = 18.7%, n = 198; bouton coverage = 10.6%, IQR = 20.7%, n = 203, p = 0.044, MW test; **Figure 1D inset**). 90% (19/198) of spines were contacted by PAPs at their membrane, compared to 84% (32/203) of boutons.

Contact near the synaptic cleft or axon-spine interface (ASI) is a critical zone for astrocytic interactions (Zheng et al., 2008; Allam et al., 2012; Rose et al., 2018). We measured the amount of astrocyte coverage at the ASI, considering both contact length and fraction of the total ASI perimeter (**Figure 1E**). Most synapses (79.8%, n = 158/198) had ASI coverage (**Figure 1F**). Of synapses with ASI coverage, the median coverage was 24.2% (n = 158) with a wide range (1.4% - 91.7%). Similar to membrane coverage, we found ASI perimeter scales strongly with PSD area (r = 0.88, p < 0.0001 **Figure 2A**). ASI coverage length and membrane coverage surface area were strongly correlated (r = 0.82, p < 0.0001; **Figure 2B**), as were the fraction of both (r = 0.76, p < 0.0001; **Figure 2C**).

**Figure 2:**
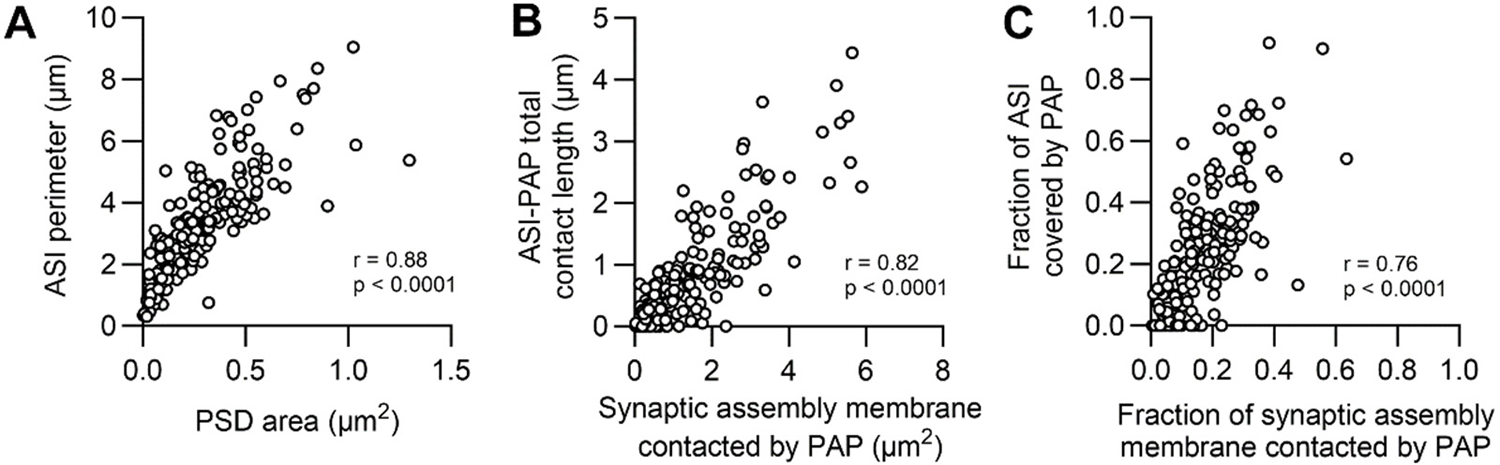
PAP coverage of the ASI covaries with synaptic assembly membrane coverage. **A.** Correlation of ASI perimeter and PSD area (r represents Spearman’s correlation coefficient). **B.** Correlation between ASI-PAP total contact length and the synaptic assembly membrane area contacted by PAP. **C**. Correlation between the fraction of ASI covered by PAP and the fraction of the synaptic assembly membrane contacted by PAP.

### PAP coverage scales with structural measures of synaptic strength

Synapse size is a correlate of synaptic strength (Holtmaat and Svoboda, 2009; Santuy et al., 2018) and the size of individual synaptic components (PSD area, spine head volume, bouton volume) varies by almost 2 orders of magnitude (Scholl et al., 2021). Thus, we investigated whether astrocytic coverage of synaptic assemblies or the ASI related to synapse structural diversity. Membrane area covered by astrocytes was positively correlated with the membrane surface area of the spine head, neck, and bouton (r = 0.54, 0.49, and 0.50, respectively; p < 0.0001; **Figure 3A-C**). In addition, the length of ASI contacted by an astrocyte was positively correlated with all structural measures (spine head volume r = 0.59; PSD area r = 0.58; bouton volume r = 0.56; ASI perimeter r = 0.62; p < 0.0001; **Figure 3D-G**). Across our measures of synaptic strength, synapses contacted by a PAP were significantly larger than those without contact (spine volume: none = 0.16 µm^2^, IQR = 0.22 µm^2^, n = 40; some = 0.35 µm^2^, IQR = 0.44 µm^2^, n = 158, p = 0.0028, MW test; PSD area: none = 0.12 µm^2^, IQR = 0.25 µm^2^, n = 40; some = 0.25 µm^2^, IQR = 0.35 µm^2^, n = 158, p = 0.0020, MW test; bouton volume: none = 0.15 µm^2^, IQR = 0.30 µm^2^, n = 40; some = 0.30 µm^2^, IQR = 0.36 µm^2^, n = 158, p = 0.0036, MW test; ASI perimeter: none = 2.29 µm, IQR = 2.15 µm; n = 41; some = 3.36 µm, IQR = 2.45 µm, n = 158, p = 0.0079, MW test; **Figure 3H-K**). While in the hippocampus and amygdala smaller spines have proportionally greater coverage (Witcher et al., 2007; Ostroff et al., 2014), our dataset from ferret visual cortex showed no significant correlations of spine head, neck, or bouton surface area to the relative fraction of PAP membrane coverage (p > 0.19), nor between spine head volume, bouton volume, PSD area, and PAP-ASI coverage fraction (p > 0.14).

**Figure 3:**
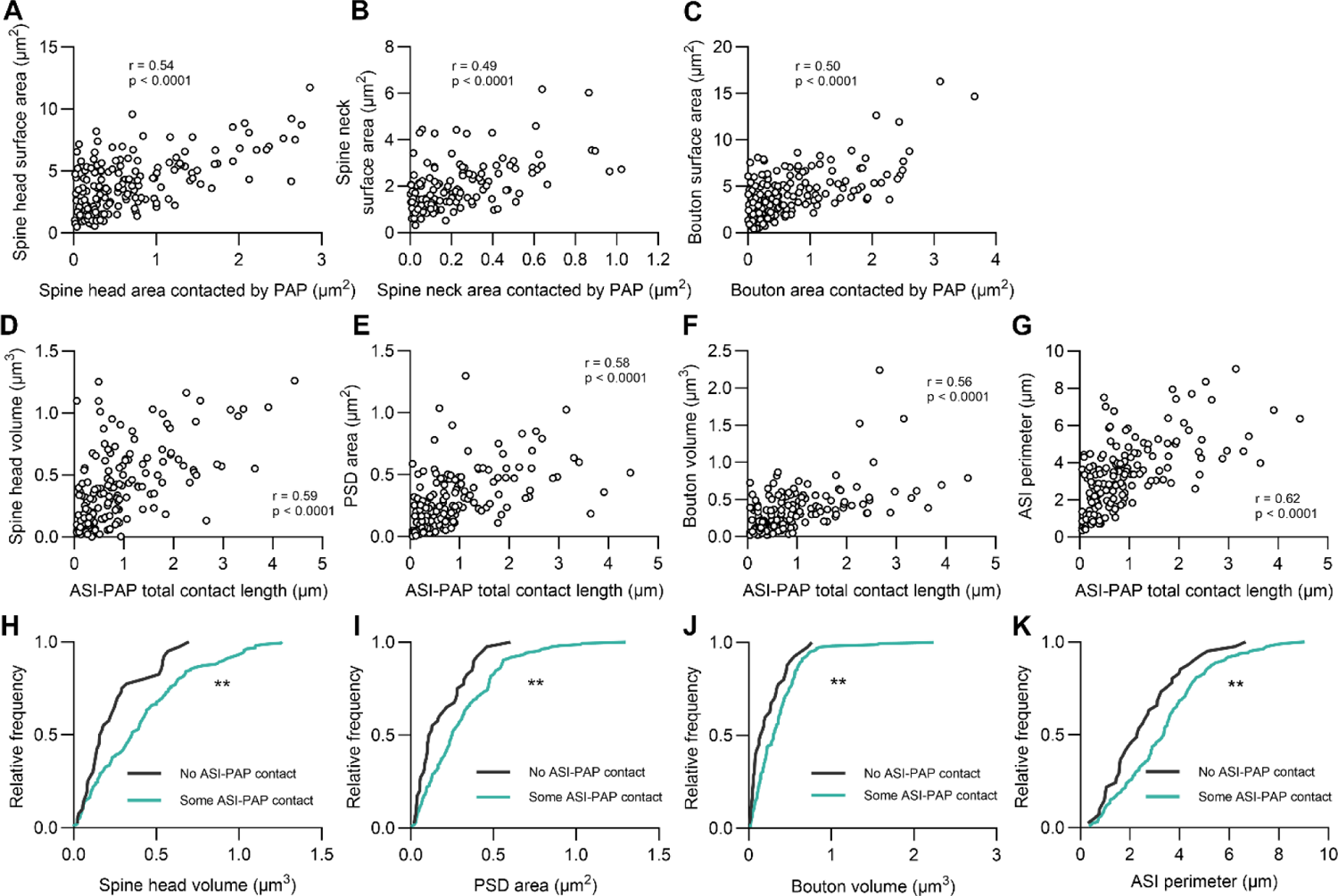
Synapse size scales with membrane and interface coverage by PAPs. **A-C.** Correlations between postsynaptic spine head, spine neck, or presynaptic bouton surface area and PAP area contacting those components (r represents Spearman’s correlation coefficient). **D-G.** Correlations between spine head volume, PSD area, bouton volume, and ASI perimeter and ASI-PAP total contact length. **H-K.** CDFs of spine head volume, PSD area, bouton volume, and ASI perimeter for synapses with no ASI-PAP contact vs. synapses with ASI-PAP contact. Statistical comparisons were made using Mann-Whitney tests. **p < 0.01.

We then categorized PSD complexity based on topology: synapses were classified as simple (macular) or complex (horseshoe, fenestrated, or segmented), following previous literature (Lushnikova et al., 2009; **Figure 4A-D**). As astrocyte coverage is higher for larger synapses and these synapses often have complex PSDs (Nieto-Sampedro et al., 1982), we examined the relationship between coverage and synapse topology. Nearly all simple and complex synapses had PAP coverage (98.5% and 99.3%, respectively). Total membrane coverage of synaptic assemblies with simple PSDs was significantly smaller than complex synapses (median coverage area: simple = 0.60 µm^2^, IQR = 1.14 µm^2^, n = 65; complex = 1.13 µm^2^, IQR = 1.89 µm^2^, n = 133, p = 0.0014, MW test; **Figure 4E**), while the fraction covered was similar (median coverage percentage: simple = 15.6%, IQR = 17.6%; complex = 13.3%, IQR = 15.0%, p = 0.45, MW test; **Figure 4F**). For ASI coverage, complex synapses had a significantly greater length of perimeter covered (median coverage length: simple = 0.265 µm, IQR = 0.635 µm; complex = 0.680 µm, IQR = 1.06 µm, p = 0.38, MW test; **Figure 4G**), but no difference in the fraction of coverage was observed (median coverage percentage: simple = 15.6%, IQR = 34.5%; complex = 20.7%, IQR = 25.8%, p = 0.38, MW test; **Figure 4H**).

**Figure 4:**
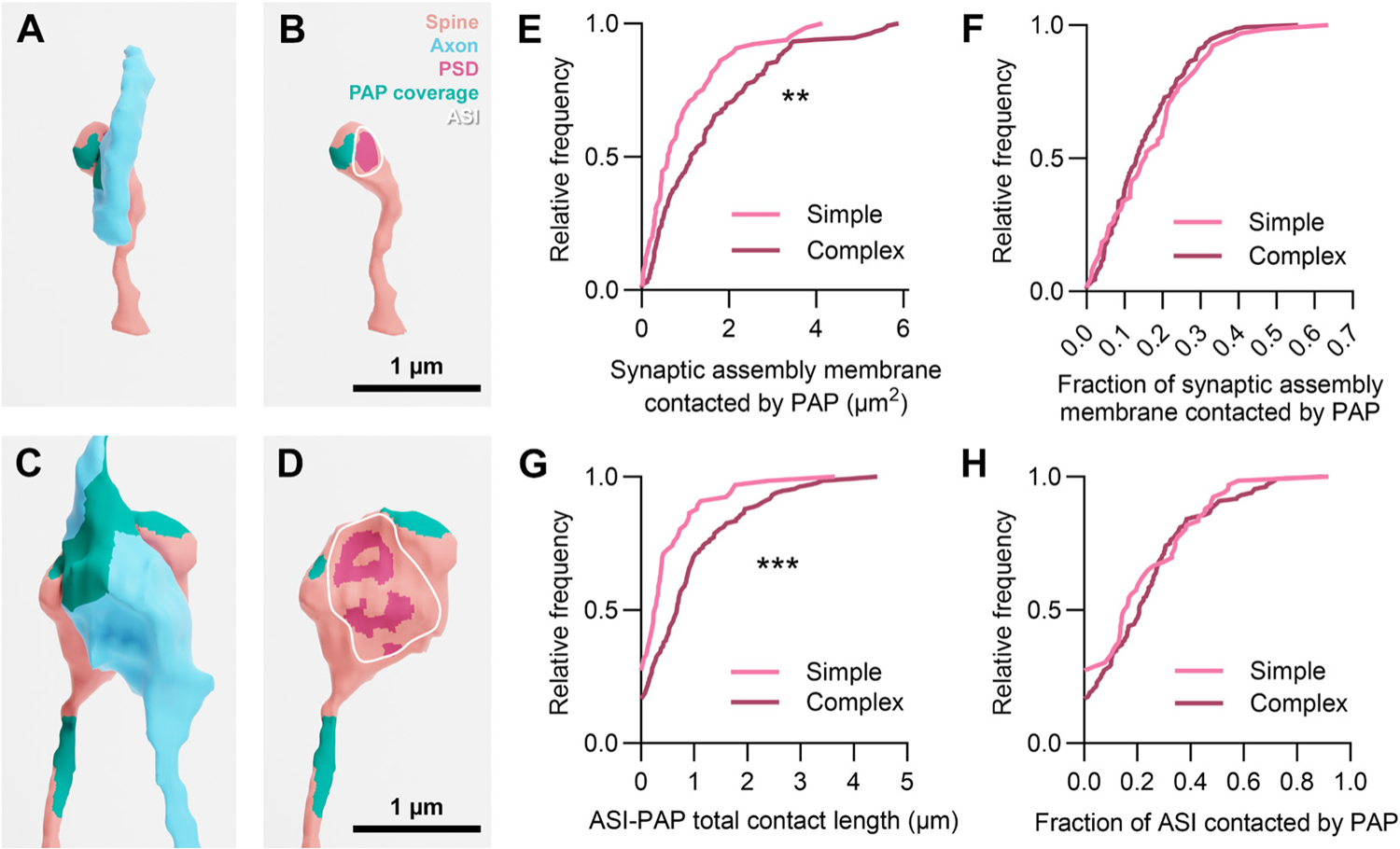
Complex synapses have greater membrane and interface coverage by PAPs than simple synapses. **A, B.** 3D EM reconstruction of a simple synapse shown with and without the presynaptic axon. **C, D.** A complex synapse shown with and without the presynaptic axon. **E.** CDF of PAP synaptic assembly coverage area, comparing spines with simple and complex synapses. **F.** CDF of the fraction of the synaptic assembly membrane that is contacted by the PAP, comparing simple and complex synapses. **G.** CDF of the ASI-PAP total contact length, comparing simple and complex synapses. **H.** CDF of the fraction of the ASI contacted by PAP, comparing simple and complex synapses. **p < 0.01, ***p < 0.001

PAPs typically made a single, continuous contact onto the ASI, but as many as 5 segments were observed on a given synapse, often spread across the interface (**Figure 5A,B**). We found that synapses with multiple segments at the ASI had a significantly larger PSD (KW test: *H* = 37.50, *df* = 3, p <0.0001; **Figure 5C**), spine head volume (KW test: *H* = 38.04, *df* = 3, p <0.0001), bouton volume (KW test: *H* = 41.12, *df* = 3, p <0.0001), and ASI perimeter (KW test: *H* = 43.61, *df* = 3, p <0.0001). Interestingly, for all four morphological measurements, size was not different between synapses without PAP contact or with only 1 contact (Post hoc Dunn’s multiple comparisons for 0 vs. 1 segment: p > 0.90; **Figure 5C**). We also observed that as the number of contacts increases, the length of those contacts increases (KW test: *H* = 42.90, *df* = 3, p < 0.0001; **Figure 5D**). Taken together, we find that the total amount (*not* relative fraction) of PAP coverage and number of discrete PAP contacts scales with the size and complexity of synaptic assemblies and ASI in layer 2/3 V1 pyramidal neurons.

**Figure 5:**
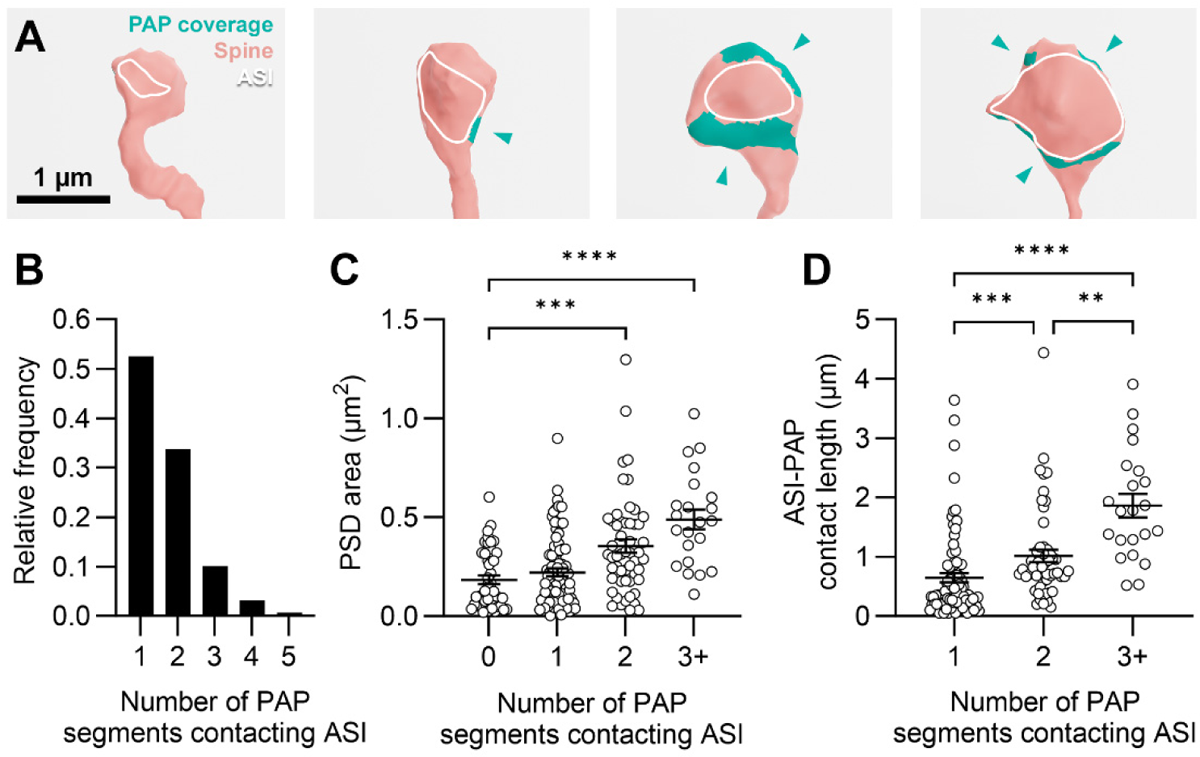
Synapses with a greater number of PAP segments contacting the ASI are larger. **A.** 3D EM reconstructions of four spines with zero, one, two, or three discrete PAP contacts at the ASI, indicated by arrows. **B.** Frequency distribution of the number of discrete PAP segments contacting the ASI. **C.** Comparison of PSD area and the number of discrete PAP segments. **D.** Comparison of segment contact length and the number of discrete PAP segments. Statistical comparisons were made using Kruskal-Wallis tests with Dunn’s multiple comparisons. **p < 0.01, ***p < 0.001, ****p < 0.0001.

### PAP coverage supports dendritic spine visual activity

Astrocyte coverage is proposed to regulate or constrain synaptic activity, but this relationship has never been examined directly (Papouin et al., 2017; Torres-Ceja and Olsen, 2022). Using our correlative dataset, we compared PAP morphology with *in vivo* two-photon Ca^2+^ imaging of dendritic spine activity evoked during visual presentation of oriented drifting gratings (**Figure 6A,B; see methods**). We then characterized the evoked Ca^2+^ responses, orientation selectivity, difference in preferred orientation between spine and somatic output, and synaptic event reliability across stimuli presented (see Methods).

**Figure 6:**
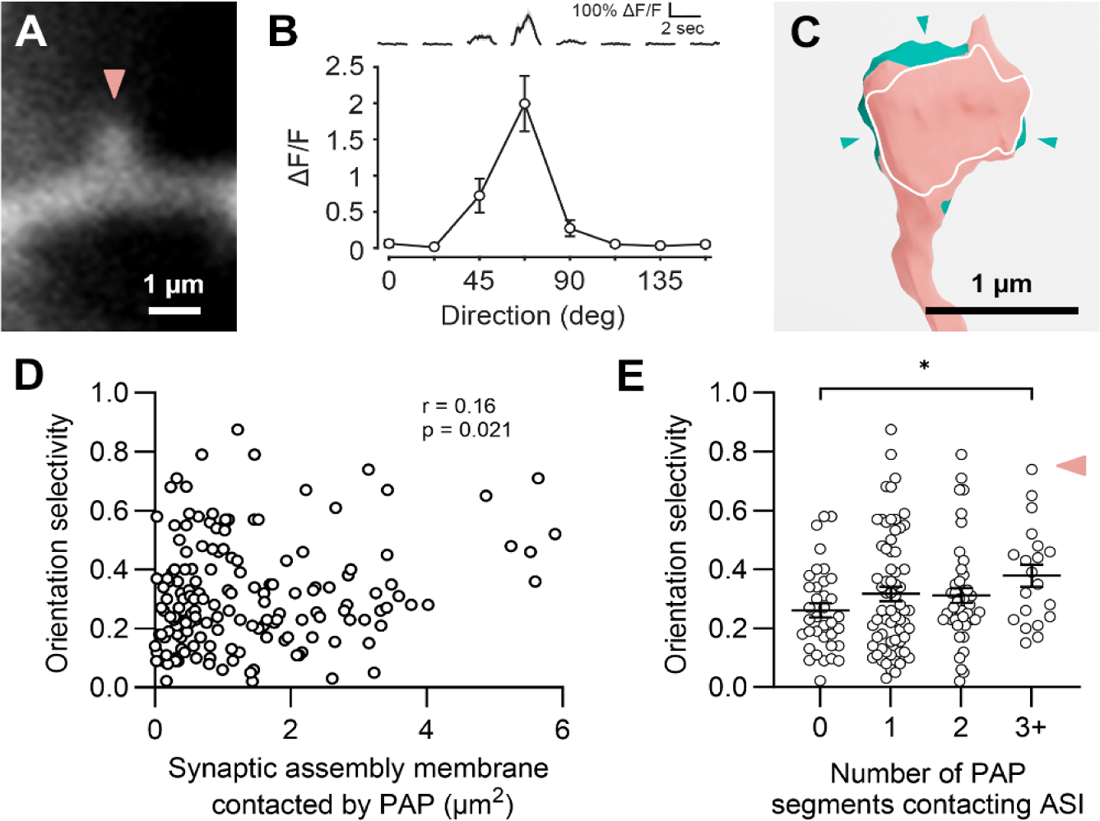
Orientation selectivity to visual stimuli weakly scales with membrane and interface coverage by PAPs. **A.** Representative image of an example spine recorded by *in vivo* 2-photon microscopy of a functionally characterized spine. **B.** Peak ΔF/F in response to visual stimuli in the preferred direction for the example spine shown in A, with a high orientation selectivity index of 0.7. **C.** 3D EM reconstruction of the same spine, showing PAP coverage (green) and 3 discrete contacts at the ASI (green arrows). **D.** Correlation of orientation selectivity and membrane coverage by PAPs (r represents Spearman’s correlation coefficient). **E.** Orientation selectivity in relation to the number of discrete PAP segments contacting the ASI. Pink arrowhead indicates the spine in A-C. *p < 0.05.

In our previous study, we discovered that stronger synapses were more selective for oriented gratings (Scholl et al., 2021). We reasoned that astrocyte coverage should also be correlated to spine selectivity. Indeed, PAP coverage of synaptic assemblies was weakly correlated to orientation selectivity (r = 0.16, p = 0.021; **Figure 6C,D**), but not for spine head coverage or bouton coverage alone (p > 0.14). ASI astrocyte contact length was also weakly correlated to orientation selectivity (r = 0.15, p = 0.048). Similar to structural measurements, relative fraction of membrane and ASI coverage were uncorrelated with spine selectivity (p = 0.44 and p = 0.19, respectively). In addition, we found that synapses with 3 or more PAP segments (**Figure 6C**) were more selective than synapse without PAP coverage (KW test: *H* = 5.95, *df* = 3, p = 0.11; Post hoc Dunn’s multiple comparisons: p = 0.048; **Figure 6E**).

Although a great number of spines are co-tuned with the somatic output (Scholl et al., 2021), there is considerable diversity in orientation preference across the population of spines on a neuron (Scholl and Fitzpatrick, 2020). We previously found that dendritic mitochondria tend to be located near spines that are more diverse in their preferences, suggesting enhanced energetic support (Thomas et al., 2023). This inspired us to consider whether differentially-tuned spines require more astrocytic support. Comparing the orientation difference between spines and parent soma to PAP coverage, we found no relationship to the total or fraction of coverage of synaptic assemblies or the ASI (p > 0.24), suggesting that astrocytes may support synaptic properties, independent of the neuronal input/output transformation.

In addition to orientation tuning preference, individual dendritic spines vary in their reliability of activation; this relates to a synapse’s efficacy and may depend on astrocytic support. To examine response reliability, we extracted stimulus-driven binary synaptic events (**Figure 7A**); defined as exponentially-filtered Ca^2+^ responses exceeding 2 standard deviations of a spine’s noise level (Scholl et al., 2021). We calculated both reliability for a spine’s preferred orientation and the average reliability across all stimuli evoking a response. Across the population, total PAP coverage of synaptic assemblies was uncorrelated with mean reliability (r = 0.070, p = 0.21, n = 129) and reliability at a spine’s preferred stimulus (r = 0.13, p = 0.067, n = 119; **Figure 7B,E**). A lack of a relationship was also observed for ASI coverage (mean reliability: r = −0.02, p = 0.40, n = 129; preferred stimulus reliability: r = 0.044, p = 0.32, n = 120).

**Figure 7:**
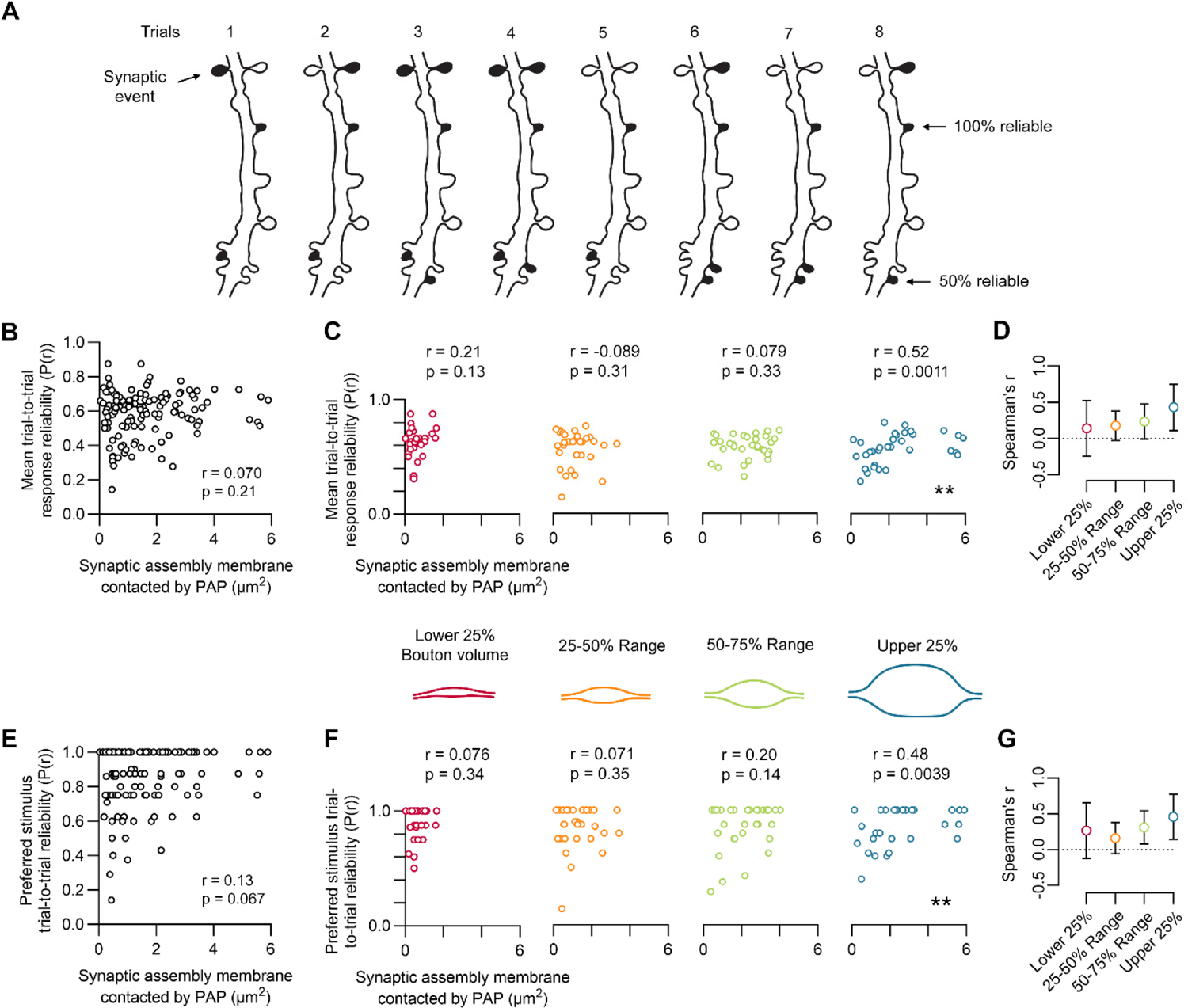
PAP coverage is correlated to synaptic reliability at strong synapses. **A.** Trial-to-trial preferred stimulus synaptic events for an example dendritic branch. **B.** Correlation of mean trial-to-trial response reliability and synaptic assembly membrane coverage by PAPs (r represents Spearman’s correlation coefficient). **C.** Subsets of points from B, split by bouton volume quartiles. **D.** Mean bootstrapped correlation coefficients (10,000 iterations) and 95% confidence intervals for each group. **E.** Correlation of preferred stimulus trial-to-trial response reliability and synaptic assembly membrane coverage by PAPs. **F.** Correlations of subsets of points from E, split by bouton volume quartiles. **G.** Mean bootstrapped correlation coefficients (10,000 iterations) and 95% confidence intervals for each group. **p < 0.01.

We next considered that such a relationship may depend on the structural properties of presynaptic boutons. Larger axonal boutons typically possess more presynaptic vesicles and have a higher likelihood of containing mitochondria for energetic support, so they may require increased astrocytic support to maintain reliable postsynaptic activation. We repeated our analyses conditioned on synapse bouton size, examining quartiles spanning the entire volume distribution (lower 25%: 0 - 0.13 µm^3^, 25 - 50%: 0.13 - 0.31 µm^3^, 50 - 75%: 0.31 - 0.53 µm^3^, upper 25%: 0.53 - 2.24 µm^3^). While synapses with smaller boutons displayed no relationship between total PAP coverage and reliability, a relationship emerged for larger boutons (**Figure 7C,F**): stimulus-driven response reliability in the largest boutons (upper 25%) was strongly correlated with astrocytic coverage (mean reliability: r = 0.52, p = 0.0011, n = 33; preferred stimulus reliability: r = 0.48, p = 0.0039, n = 30). The same relationships were found for the largest 25% of dendritic spine heads (mean reliability: r = 0.44, p = 0.0056, n = 32; preferred stimulus reliability: r = 0.40, p = 0.0015, n = 30). Although only synapses with the largest boutons displayed a statistically significant correlation between reliability and bouton size, we found that the correlation magnitude, calculated by a bootstrap analysis of correlation between PAP coverage and reliability, steadily increased across quartiles (**Figure 7D,G**). As these data suggest that visual response reliability of large synapses is constrained by the amount of astrocytic coverage, it stands to reason that the same correlation is evident when comparing coverage fraction. As expected, reliability was correlated to the fraction of PAP coverage in synapses with the largest boutons (mean reliability: r = 0.53, p = 0.0007, n = 33; preferred stimulus reliability: r = 0.47, p = 0.0048, n = 30) and the fraction of coverage along the ASI (mean reliability: r = 0.33, p = 0.029, n = 33; preferred stimulus reliability: r = 0.38, p = 0.021, n = 30).

Altogether, our data provide two findings regarding dendritic spine Ca^2+^ activity during visual stimulation. First, spine selectivity is correlated with astrocyte coverage, similar to spine size and structural complexity. Second, the reliability of visually-evoked synaptic events correlates with the amount or degree of PAP coverage in the strongest synapses, suggesting that the glutamate spillover and vesicular content of these synapses may require astrocytic support for high-fidelity signaling.

## Discussion

We examined two fundamental questions about the structural relationship between astrocytes and synapses: (1) can tripartite synapse structure be described by a universal rule and (2) does tripartite synapse structure relate to synaptic functional activity? Using volume EM 3D reconstructions of layer 2/3 pyramidal neurons from ferret primary visual cortex, we measured the coverage of perisynaptic astrocytic processes (PAP) over the synaptic assembly membrane as well as coverage at the axon-spine interface (ASI) perimeter. We found nearly all synapses had coverage at the synaptic assembly and the vast majority had coverage at the ASI. We observed that synapse size and complexity scale positively with membrane and ASI coverage by PAPs, while the fraction of coverage is uncorrelated. For functional response properties, we found that synapses with a high degree of coverage are more selective to visual stimuli, but there was no difference between synapses co-tuned and differentially-tuned compared to the somatic output. We also observed that for strong synapses, the trial-to-trial reliability of visually-evoked responses depended on the amount or fraction of astrocytic coverage over the synaptic assembly and the ASI. Our study is the first to link astrocyte coverage to both structure and function of individual synapses and contributes to building evidence that different brain regions exhibit specialization in the formation of tripartite synapses.

### Covariation in PAP coverage and synapse size may reflect sensory cortex specialization

The structural relationship between PAP coverage and synapse size has been previously investigated, with different conclusions being drawn depending on the brain region under study (Genoud et al., 2006; Witcher et al., 2007; Medvedev et al., 2014; Herde et al., 2020). In both the hippocampus and lateral amygdala, ASI coverage length by astrocytes and PSD area exhibited no correlation (Witcher et al., 2007; Ostroff et al., 2014), while in the barrel cortex, there is a strong relationship, similar to our own findings (Genoud et al., 2006). These differences could reflect regional variability or species specialization of PAP-synapse interactions. For example, in the human cortex, astrocytes are highly complex and cover vastly more synapses than in mouse, leading to the idea that astrocyte proliferation may promote ‘increased functional competence’ (Bushong et al., 2002; Oberheim et al., 2009). Similarly, ferret cortical astrocytes are larger than those observed in mouse cortex (López-Hidalgo et al., 2016), potentially supporting greater functional demands of their cortical circuits. This may, in part, be due to the functional demands of a dedicated sensory cortical circuit such as the primary visual cortex; ferrets are a highly visual animal and the primary visual cortex encompasses a significant portion of the cortical surface. In line with this idea, rodent barrel cortex, a primary sensory area crucial for processing information from the whiskers (Diamond et al., 2008), also displays a high degree of PAP coverage and a correlation between PAP coverage and PSD size (Genoud et al., 2006). Together, these studies suggest specialized astrocytic interactions in sensory cortical circuits as compared to subcortical brain areas such as the hippocampus and amygdala.

### PAP coverage and the limitation of glutamate spillover

Glutamate spillover from the synaptic cleft is proposed to be regulated by astrocytic contacts via the activity of Na^+^-dependent glutamate transporters (Henneberger et al., 2020; Herde et al., 2020). In CA1 stratum radiatum, where coverage and synapse size are not correlated, expansion microscopy revealed that small spines have proportionally greater access to the glutamate transporter GLT1 (Herde et al., 2020), leading to less spillover as compared to larger synapses and insulation from plasticity induced by extrasynaptic activation of NMDA receptors (Liu et al., 2004; Zheng et al., 2008; Papouin et al., 2012; Herde et al., 2020). Our data, on the other hand, reveal no coverage biases between small and large synapses, allowing for relatively similar insulation from glutamate spillover and isolation from extrasynaptic-induced plasticity. Whether this reflects a regional specialization or species difference is still unclear. We hypothesize that the high degree of PAP coverage may provide a computational advantage to networks through increased synaptic independence and isolation of signals (Scimemi et al., 2004; Verkhratsky and Nedergaard, 2018).

Glutamate spillover is likely not only controlled by the amount of astrocytic coverage, but also the number and distribution of contacts. Nearly half of synapses in our dataset had two or more contacts, often distributed around the perimeter of the ASI (**Figure 5**). Interestingly, while we found synapses with at least some PAP coverage were stronger, synapses with only one contact at the ASI were no stronger than those with none at all. It is reasonable to expect that a greater number of contacts create a more uniform distribution of glutamate transporters around the ASI, reducing unidirectional spillover and assisting in glutamate uptake to support stronger synapses. In support of this idea, induction of LTP in hippocampal organotypic slices increases the number of discrete contacts on dendritic spines (Lushnikova et al., 2009). Simulations of glutamate uptake with varying configurations of astrocyte segments at the ASI may be helpful to further illustrate why this system appears to be preferred at stronger synapses.

### PAP coverage and synaptic reliability

We were surprised to discover that response reliability in strong, but not weak synapses, was correlated with astrocytic coverage (**Figure 7**). These data suggest that larger synapses, with greater numbers of synaptic vesicles, require more efficient turnover of excitatory neurotransmitter to facilitate high fidelity activation. Astrocytes take up and convert glutamate into glutamine, which is then released to the extracellular space, taken up by the presynaptic bouton, and converted back into glutamate (Norenberg and Martinez-Hernandez, 1979; Hertz et al., 1999; Bak et al., 2006; Cheung et al., 2022). This cycle is critical for the maintenance of cytoplasmic glutamate levels, synaptic transmission, sensory processing, and plasticity (Conti and Minelli, 1994; Masson et al., 2006; Holten and Gundersen, 2008; Gaisler-Salomon et al., 2012; Cheung et al., 2022). Interestingly, small synapses frequently displayed reliable activation by visual stimuli, even with little astrocytic coverage. A possible explanation for this is that presynaptic glutamate transporters (e.g. GLT-1a) contribute to neurotransmitter reuptake when astrocyte coverage is sparse, but little is known about the physiological significance of these transporters in brain tissue (Chen et al., 2004; Rimmele and Rosenberg, 2016). Another possibility is that response reliability depends on other factors such as the presence of a spine apparatus, postsynaptic receptor density, or presynaptic vesicle release probability.

### Limitations and future directions

While our work provides several new insights into the relationship between synapse structure-function and astrocyte coverage, there are several limitations. We do not take into account astrocytic glutamate transporters at the membrane, which can modulate synaptic transmission following plasticity and adjust their distribution based on neuronal activity (Zhou and Sutherland, 2004; Murphy-Royal et al., 2015; Al Awabdh et al., 2016). Additionally, the abundance of presynaptic glutamine transporters likely contributes to the glutamate replenishment cycle and maintenance of synaptic transmission (Marx et al., 2015). In brain regions with a high degree of astrocyte proliferation, glutamate/glutamine cycling may be tightly modulated through the control of these transporters, resulting in further synaptic isolation and increased reliability.

Another limitation of our structural analysis, common among EM studies using chemical fixation and dehydration, is that the extracellular space (ECS) is not well preserved. Tissue shrinkage due to changes in osmotic pressure induced by conventional EM preparation results in ECS collapse, obfuscating the study of neurotransmitter spillover and uptake (Korogod et al., 2015). Recent advancements in sample preparation have led to preservation of the osmotic balance, through the addition of the sugar mannitol to the fixation solution (Karlupia et al., 2023; Lu et al., 2023). Preservation of the ECS will serve as a critical step forward for the study of astrocyte support.

Despite the structure-function relationships between synapses and astrocytes observed by many studies, particularly those done in the hippocampus, few relationships are likely to be universal. Considering the variability in astrocyte morphology and coverage, it is clear that there is a need for more comparative studies. In addition, *in vivo* recordings of astrocyte function and spine activity may expand our understanding of tripartite synapse dynamics. While we investigated spine activity in response to visual stimuli, our CLEM data represent a single snapshot of astrocyte-synapse interaction. Both spines and astrocytes are highly dynamic structures, changing over time and in response to sensory activity (Genoud et al., 2006; Bernardinelli et al., 2014b; Henneberger et al., 2020; Steffens et al., 2021). Thus, the investigation of *in vivo* spine and astrocyte activity, particularly using longitudinal imaging, super-resolution microscopy, and correlative light and electron microscopy, will be critical to better understand the interactions within tripartite synapses and their dynamics across brain regions and model systems.

## Acknowledgements

The authors thank Nicole Shultz and Rachel Satterfield for their help with perfusions and fixative preparation, Skylar Anthony for her assistance with astrocyte measurements, the Fitzpatrick lab for useful discussions, and the MPFI ARC for animal care. The authors thank the GENIE project for access to GCaMP6.

